# The effect of corticosterone on the acquisition of Pavlovian conditioned approach behavior is dependent on sex and vendor

**DOI:** 10.1101/2024.03.20.586009

**Authors:** Alexandra Turfe, Sara R. Westbrook, Sofia A. Lopez, Stephen E. Chang, Shelly B. Flagel

**Affiliations:** Michigan Neuroscience Institute, University of Michigan; Neuroscience Graduate Program, University of Michigan; Department of Psychiatry, University of Michigan, Ann Arbor, 48109

**Keywords:** corticosterone, Pavlovian conditioning, reward, sign-tracking, rats

## Abstract

Cues in the environment become predictors of biologically relevant stimuli, such as food, through associative learning. These cues can not only act as predictors but can also be attributed with incentive motivational value and gain control over behavior. When a cue is imbued with incentive salience, it attains the ability to elicit maladaptive behaviors characteristic of psychopathology. We can capture the propensity to attribute incentive salience to a reward cue in rats using a Pavlovian conditioned approach paradigm, in which the presentation of a discrete lever-cue is followed by the delivery of a food reward. Upon learning the cue-reward relationship, some rats, termed sign-trackers, develop a conditioned response directed towards the lever-cue; whereas others, termed goal-trackers, approach the food cup upon lever-cue presentation. Here, we assessed the effects of systemic corticosterone (CORT) on the acquisition and expression of sign- and goal-tracking behaviors in male and female rats, while examining the role of the vendor (Charles River or Taconic) from which the rats originated in these effects. Male and female rats from Charles River had a greater tendency to sign-track than those from Taconic. Administration of CORT enhanced the acquisition of sign-tracking behavior in males from Charles River and females from both vendors. Conversely, administration of CORT had no effect on the expression of the conditioned response. These findings demonstrate a role for CORT in cue-reward learning and suggest that inherent tendencies towards sign- or goal-tracking may interact with this physiological mediator of motivated behavior.

**Highlights:**

- Male and female rats from Charles River exhibit more sign-tracking relative to those from Taconic.
- Corticosterone increases the acquisition of sign-tracking in male rats from Charles River.
- Corticosterone increases the acquisition of sign-tracking in female rats, regardless of vendor.
- There is no effect of corticosterone on the expression of sign-tracking behavior in either male or female rats.

## Introduction

In our daily lives we are exposed to an abundance of stimuli, or cues, in our environment that guide our choices and actions. For example, the smell of French fries or sight of a fast-food sign may elicit craving and cause us to eat even if we are not hungry. The ability of such cues to guide behavior occurs through Pavlovian or associative learning based on prior experiences (Rescorla, 1988). Previously neutral cues gain predictive value and influence behavior in an adaptive way. However, in addition to predictive value, these cues can also gain incentive value through the process of incentive salience attribution (Robinson & Berridge, 1993). This process enables cues to act as powerful “lures” that can trigger maladaptive behaviors like overconsumption of food or relapse to drug-use.

Individuals vary in the way in which they learn about and respond to cues in the environment, and we can capture this individual variation using an animal model. When rats are exposed to a Pavlovian conditioned approach (PavCA) paradigm in which a discrete lever-cue (conditioned stimulus, CS) is presented, and immediately followed by the delivery of a food reward (unconditioned stimulus, US), distinct conditioned responses emerge (Boakes, 1977; Robinson & Flagel, 2009). Some rats, sign-trackers (STs), direct their behavior towards the lever-cue. Whereas others, goal-trackers (GTs), interact with the location of impending reward delivery, or the food cup. While both sign-trackers and goal-trackers attribute predictive value to the lever-cue, sign-trackers also attribute incentive motivational value to the lever-cue, which renders the cue itself attractive and desirable (Saunders & Robinson, 2010, 2011; Yager et al. 2015; Yager & Robinson, 2013). Thus, the PavCA paradigm allows us to experimentally parse predictive vs. incentive learning styles and to uncover the underlying neural and biological mechanisms.

The sign-tracker/goal-tracker animal model revealed that ventral tegmental area dopamine neuron activity (Iglesias et al., 2023) and dopamine release in the nucleus accumbens (NAc) (Flagel, Clark et al. 2011) is necessary for the attribution of incentive value to reward cues, but not predictive value. Dopamine in the NAc also increases in response to stressors (Kalivas & Duffy, 1995) and following the administration of glucocorticoids (Piazza et al. 1996). Glucocorticoids (cortisol in humans; corticosterone (CORT) in rodents) are known traditionally for their role in mediating the stress response and regulating hypothalamic-pituitary-adrenal (HPA) axis activity via their actions at glucocorticoid receptors (Henry, 1992). Importantly, however, exposure to rewards (e.g., food, sex, drugs) can also elicit a rise in corticosterone (Piazza & Le Moal, 1997). Further, this rise in corticosterone can itself be reinforcing, as rats will intravenously self-administer corticosterone at levels similar to the physiological stress response (Piazza et al., 1993). Glucocorticoids have also been shown to facilitate the reinforcing effects of cocaine such that elimination of endogenous corticosterone via adrenalectomy results in a significant downward shift in a self-administration dose-response curve, and this effect is reversed with administration of CORT (Deroche et al., 1997). Together, these findings led to the notion that glucocorticoids are one of the biological factors underlying vulnerability to substance use and abuse (Piazza & Le Moal, 1996).

The propensity to attribute incentive salience to reward cues has also been related to glucocorticoids. A single PavCA session elicits a rise in CORT to a greater extent in male rats that become sign-trackers relative to those who become goal-trackers (Flagel et al., 2009). Additionally, systemic administration of a GR antagonist in male Japanese quail decreases sign-tracking behavior in a dose-dependent manner (Rice et al., 2018; Rice et al., 2019). These findings suggest that CORT facilitates the attribution of incentive motivational value to reward cues, and, in turn, sign-tracking behavior. To determine if this is indeed the case, we assessed the effects of CORT administration on both the acquisition and expression of Pavlovian conditioned approach behavior. We found that the administration of systemic CORT did not impact the expression of Pavlovian conditioned approach behavior but did affect the acquisition of sign-tracking behavior. Importantly, this effect was dependent on the sex of the rat and the vendor from which the rats originated.

## Materials and Methods

This study involved three separate experiments, as outlined in Figure 1. Experiment 1 consisted of male rats, Experiment 2 consisted of female rats, and Experiment 3 had both male and female rats. Experiments 1 and 2 examined the effects of CORT on the acquisition of Pavlovian conditioned approach behavior and were identical in protocol, except for vaginal lavages performed in Experiment 2; while Experiment 3 examined the effects of CORT on the expression of Pavlovian conditioned approach behavior.

**Figure 1.**
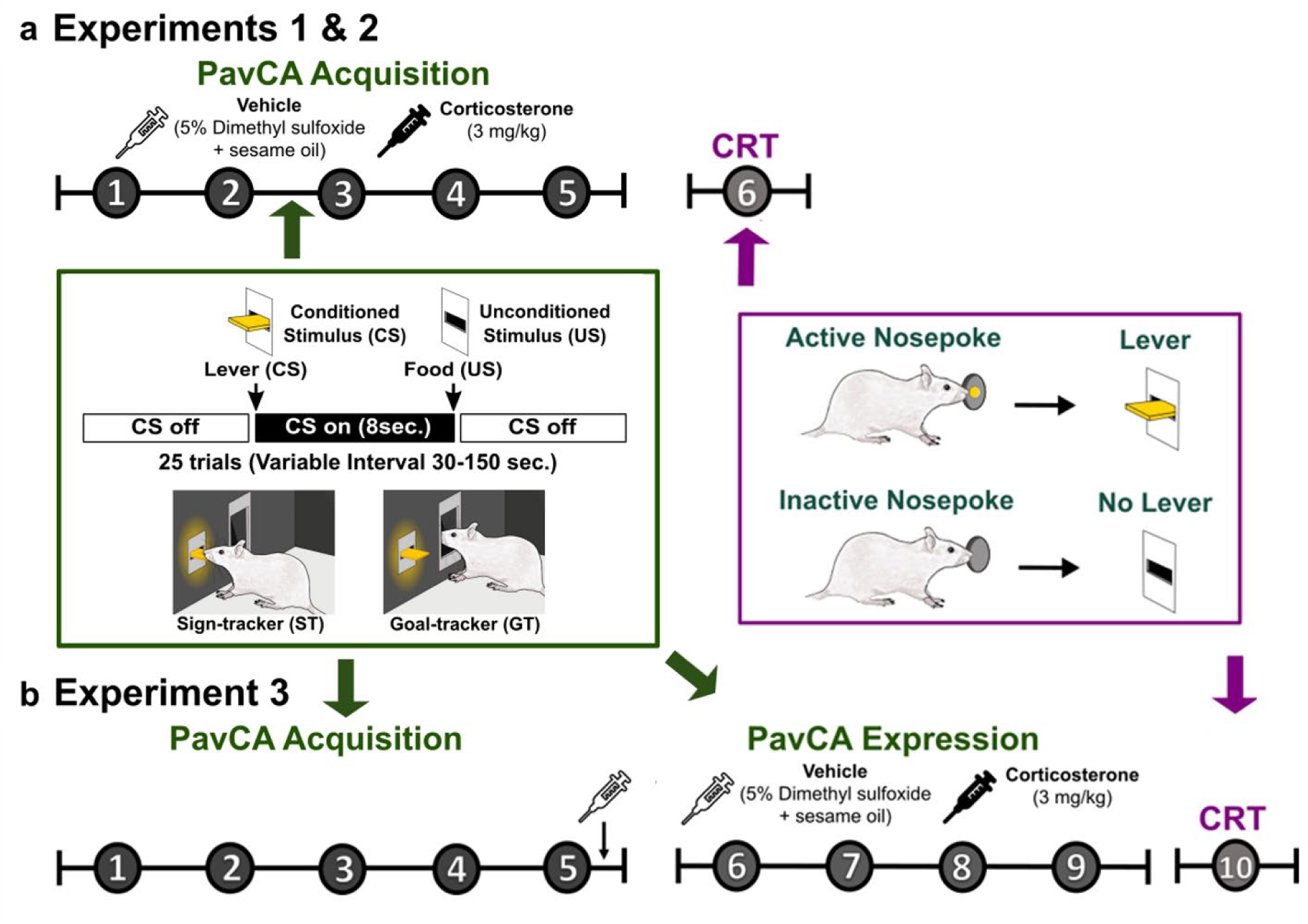
Experimental timeline. (a) Experimental timeline for Experiments 1 and 2. Schematic illustrating Pavlovian Conditioned Approach (PavCA) shown in green and the Conditioned Reinforcement Test (CRT) shown in purple. Rats received either vehicle or corticosterone (3 mg/kg) prior to each of the first 5 PavCA sessions for Experiments 1 and 2. (b) Experimental timeline for Experiment 3, during which rats received either vehicle or corticosterone (3 mg/kg) prior to sessions 6-9 of PavCA.

### Subjects

Male and female Sprague-Dawley rats were obtained from different vendors, as indicated below for each experiment. All rats had *ad libitum* access to food and water. Upon arrival, rats were pair-housed with the same sex in a housing room with a 12-hr light/dark cycle with lights on at 07:00. Rats were left undisturbed to acclimate to the housing conditions for at least seven days after arrival. All experimental procedures took place during the light phase of the cycle (between 09:00 and 14:00). Guidelines put forth in The Guide for the Care of and Use of Laboratory Animals: Eighth Edition (2011, National Academy of Sciences) were followed, and all experimental procedures were approved by the University of Michigan Institutional Animal Care and Use Committee.

#### Experiment 1

A total of 54 male Sprague-Dawley rats, weighing 225-275g upon arrival, were used. 22 rats were from Taconic Biosciences (colony IBU16, Cambridge City, IN, USA) and 32 rats were from Charles River Breeding Labs (colony R04, Raleigh, NC, USA). Two rounds of data collection occurred, with the first round consisting of 24 rats and the second consisting of 30 rats. Each round was conducted by two different experimenters, but both rounds of rats underwent identical experimental procedures, including behavioral testing for PavCA and conditioned reinforcement (CRT). Three rats were excluded from all PavCA and CRT analyses due to equipment malfunction (n=2 VEH Taconic, n=1 CORT Taconic). For two other rats (n=1 VEH Taconic; n=1 CORT Taconic), only PavCA session 5 data was excluded due to experimenter error. Conditioned reinforcement test data was excluded for all measures for two additional rats—one (n=1 CORT Charles River) due to equipment malfunction and one (n=1 CORT Charles River) was identified as a significant outlier with a z-score ≥ 5 for at least one of the measures. Statistical analyses were conducted on the remaining rats (Table 1).

**Table 1.** Sample sizes for statistical analyses for each experiment. (a) Final sample sizes for analyses of data collected during Pavlovian conditioned approach (PavCA). (b) Final sample sizes for analyses of data collected during the conditioned reinforcement test (CRT).

|  | **a** | **Sample Sizes for PavCA analyses** |  |  |  |  |  |  |  |
| --- | --- | --- | --- | --- | --- | --- | --- | --- | --- |
|  |  | Experiment 1<br>(Males) |  | Experiment 2<br>(Females) |  | Experiment 3<br>(both sexes) |  |  |  |
|  |  | VEH<br>Males | CORT<br>Males | VEH<br>Females | CORT<br>Females | VEH<br>Males | CORT<br>Males | VEH<br>Females | CORT<br>Females |
| Taconic |  | 9 | 10 | 15 | 15 | 28 | 23 | 23 | 29 |
| Charles River |  | 16 | 16 | 14 | 14 | 13 | 18 | 18 | 14 |
|  | **b** | **Sample Sizes for CRT analyses** |  |  |  |  |  |  |  |
|  |  | Experiment 1<br>(Males) |  | Experiment 2<br>(Females) |  | Experiment 3<br>(both sexes) |  |  |  |
|  |  | VEH<br>Males | CORT<br>Males | VEH<br>Females | CORT<br>Females | VEH<br>Males | CORT<br>Males | VEH<br>Females | CORT<br>Females |
| Taconic |  | 9 | 10 | 15 | 15 | 27 | 22 | 22 | 29 |
| Charles River |  | 16 | 14 | 14 | 14 | 13 | 18 | 16 | 13 |

#### Experiment 2

A total of 60 female Sprague-Dawley rats, weighing 200-300g upon arrival, were used. Two rounds of data collection occurred with the same experimenters for both. The first round consisted of 30 rats from Taconic Breeding Labs (colony IBU16, Cambridge City, IN, USA), while the second round consisted of 30 rats from Charles River Breeding Labs (colony R04 Raleigh, NC, USA). Two rats were excluded from all PavCA and CRT analyses due to equipment malfunction (n=2 CORT Charles River). Statistical analyses were conducted on the remaining rats (Table 1).

#### Experiment 3

Both male and female rats were used, and the study was conducted in three rounds with identical procedures but different experimenters. 84 male Sprague-Dawley rats, weighing 225-275g upon arrival, were used. 52 male rats were from Taconic Biosciences (colony IBU16, Cambridge City, IN, USA), and 32 male rats were from Charles River Breeding Labs (colonies R04 and R08 Raleigh, NC, USA). 84 female Sprague-Dawley rats, weighing 200-300g upon arrival, were used. 52 female rats were from Taconic Biosciences (colony IBU16, Cambridge City, IN, USA), and 32 female rats were from Charles River Breeding Labs (colonies R04 and R08 Raleigh, NC, USA). The first round consisted of 48 rats (24 males and 24 females), the second round consisted of 60 rats (30 males and 30 females), and the third round consisted of 60 rats (30 males and 30 females). Two rats were excluded from all analyses—one rat (n=1 Male VEH Charles River) died on the second day of injections in the experiment, and another rat (n=1 Male VEH Taconic) did not consume all of the pellets delivered during the PavCA sessions. Data for all measures during CRT were excluded for one rat due to equipment malfunction (n=1 F VEH Taconic) and five rats that were identified as significant outliers with a z-score ≥ 5 on at least one measure (n=1 F CORT Charles River, n= 2 F VEH Charles River, n=1 M VEH Taconic, n=1 M CORT Taconic). Statistical analyses were conducted on the remaining rats (Table 1).

### Pavlovian conditioned approach (PavCA)

All PavCA pre-training and training took place in standard behavioral testing chambers (MED Associates, St. Albans, VT, USA; 20.5 x 24.1 cm floor area, 29.2 cm high) located inside a room with red lighting. Each chamber was located inside a sound-attenuating cabinet with a ventilation fan that provided constant air circulation and white noise. A food-cup was centered on one of the chamber walls and placed 6 cm above the grid floor. The food-cup contained an infrared beam, and each time this beam was broken a magazine entry was recorded as a “contact”. A retractable lever, which illuminated upon presentation, was placed 6 cm above the grid floor on either the right or left of the food-cup with location counterbalanced across rats. Each time a force of at least 10 g was placed upon the lever-cue, a contact was recorded. A white house light was placed 1 cm from the top of the chamber on the wall opposite the lever and food-cup. The house light was illuminated at the beginning of the session and remained on for the duration of the session.

Rats first underwent brief handling by the experimenter(s) for two consecutive days, during which they were given approximately 25 banana-flavored food pellets (Bio-Serv, Flemington, NJ, USA) to familiarize them with the reward that would be used during PavCA training. After the two days of handling, rats underwent a single pre-training session. Five minutes after being placed in the chamber, the house light was illuminated and the pre-training session commenced, which lasted approximately 12.5 minutes. The pre-training session consisted of 25 trials during which the lever remained retracted, and pellets were delivered on a variable interval 30-s schedule (VI 30, range 0-60 s).

The day following the pre-training session, rats in Experiments 1 and 2 underwent a total of five consecutive PavCA training sessions as shown in the timeline (Figure 1a). Rats in Experiment 3 underwent a total of nine consecutive PavCA training sessions, in which sessions 1-5 were deemed the “PavCA Acquisition” phase of the experiment, and sessions 6-9 were the “PavCA Expression” phase of the experiment (Figure 1b). During the PavCA sessions, the illuminated lever-cue (conditioned stimulus, CS) was presented in the chamber for 8 seconds before being retracted. Lever-cue retraction was immediately followed by delivery of the food pellet reward (unconditioned stimulus, US) into the food-cup. Each session consisted of 25 trials on a variable interval 90 s schedule (VI 90, range 30-150s) and lasted approximately 40 minutes. All rats underwent one daily training session for 5 (Experiments 1 and 2) or 9 consecutive days (Experiment 3) between the hours of 10:00-12:00.

The following behavioral measures were recorded by Med Associates software during each PavCA session: (1) number of lever-cue contacts, (2) latency to first lever-cue contact, (3) probability to approach the lever-cue, (4) number of food-cup contacts during presentation of the lever-cue, (5) latency to first food-cup contact during presentation of the lever-cue, (6) probability of approaching the food-cup during presentation of the lever-cue, and (7) number of food-cup contacts during the inter-trial interval. These values were then used to calculate three measures of approach behavior that comprise the PavCA index: (1) response bias = [(total lever-cue contacts – total food-cup contacts) ÷ (total lever-cue contacts + total food-cup contacts)], (2) probability difference = [probability to approach the lever-cue – the probability to approach the food-cup], (3) latency difference = [± (latency to contact the lever-cue – latency to contact the food-cup) ÷ 8]. A PavCA index score was calculated for each session using the formula: [(response bias + probability difference + latency difference) ÷ 3] as initially described in Meyer et al., (2012). PavCA index scores ranged from -1 to +1 with a more positive score indicating a greater propensity to contact the lever-cue (i.e., sign-tracking) and a more negative score indicating a greater propensity to contact the location of reward delivery (i.e., goal-tracking).

### Conditioned Reinforcement Test (CRT)

Following the fifth session of PavCA training, all rats in Experiments 1 and 2 underwent a conditioned reinforcement test. Rats in Experiment 3 underwent testing for conditioned reinforcement following the ninth session of PavCA training (data not shown). The purpose of the CRT was to assess the effect of *prior* CORT administration on the conditioned reinforcing properties of the lever-cue; thus, CORT was not administered immediately prior to the test. The testing chambers were reconfigured such that the lever replaced the food-cup in the center of the wall. Two nose ports were placed on either side of the lever, with one acting as the “active” port, in which nose pokes resulted in a 2-sec presentation of the illuminated lever, and the other acting as the “inactive” port, in which responses were recorded but without consequence. To minimize side bias, the nose port placed opposite to the lever’s original position was designated the “active” port, and the nose port placed in the lever’s original position was designated the “inactive” port. One minute after rats were placed into the chamber, the house light turned on and remained on for the duration of the session. During the session, the number of pokes into the active and inactive nose ports and the number of lever contacts were recorded. A composite incentive value index (Hughson et al., 2019) was calculated using this formula: [(active pokes + lever contacts) – (inactive pokes)]. The duration of the conditioned reinforcement session was 40 min.

### Pharmacological treatment

In Experiments 1 and 2, immediately after the pre-training session, all rats (regardless of experimental group) were administered an intraperitoneal (i.p.) injection of vehicle solution (VEH, 5% dimethylsulfoxide (DMSO) in sesame oil) to familiarize them with the injection procedure. Rats were assigned to either a corticosterone (CORT; Lkt Laboratories) or a VEH group, counterbalanced across vendor. Thirty minutes prior to each daily PavCA session (1-5), rats received an i.p. injection of VEH or CORT (3 mg/kg, i.p.).

In Experiment 3, immediately after the 5^th^ PavCA session, all rats (regardless of experimental group) were administered an i.p. injection of VEH in order to familiarize them with the injection procedure. Rats were assigned to either a CORT or VEH group counterbalanced across sex and vendor. Additionally, treatment assignments were counterbalanced by average PavCA index on sessions 4 and 5, such that the treatments were equally split within goal-trackers (PavCA index ≤ -0.45), intermediated responders (-0.45 < PavCA index < +0.45), and sign-trackers (PavCA index ≥ +0.45). Thirty minutes prior to each daily PavCA session (6-9), rats received an i.p. injection of VEH or CORT (3 mg/kg, i.p.).

### Monitoring the estrous cycle

Experiments 2 and 3 included female rats that were weighed and monitored daily (15:00-18:00) for their estrous cycle by vaginal lavages as previously described (Becker et al., 2005). Lavages began on the first day of handling and continued to be performed daily, after each session, for the duration of the experiment. Data are not included due to small sample sizes after assessing the data by day of experiment and estrous cycle phase.

### Statistical Analysis

The Statistical Package for the Social Sciences (SPSS) program version 27.0 (IBM, Armok, NY, USA) was used to analyze behavioral outcome measures. When session was included as a factor, linear mixed-effects models were conducted, and the model with the best covariance structure for each dataset was selected using the lowest Akaike’s information criterion. Statistical significance was set at p ≤ 0.05. When significant main effects or interactions were detected, Bonferroni *post hoc* comparisons were performed. All figures were made using GraphPad Prism 10.

Since Experiments 1 and 2 were conducted separately, behavior across PavCA sessions was analyzed independently for each sex using linear mixed effects models with lever-cue- and food-cup-directed behaviors (i.e., contacts, probability, latency) as dependent variables (Figures 2 and 5) and treatment and vendor as between-subject factors. In addition, the average PavCA index from sessions 4 and 5 was used as a dependent variable and compared between treatment and vendors (Figures 3a and 6a) using a two-way ANOVA. The effect of treatment and vendor on PavCA index across sessions was also assessed (Figures 3b-c and 6b-c) using linear mixed-effects models. For Experiment 3, males and females were run concurrently, thus PavCA index (Figure 8a) was analyzed using linear mixed-effects models with session as a within-subjects factor and treatment, vendor, and sex as between-subjects factors. In addition, the average PavCA index from sessions 4 and 5 (acquisition phase) and sessions 6-9 (expression phase) were compared between treatment, vendors, and sex (Figures 8b-c) using a repeated measures ANOVA.

**Figure 2.**
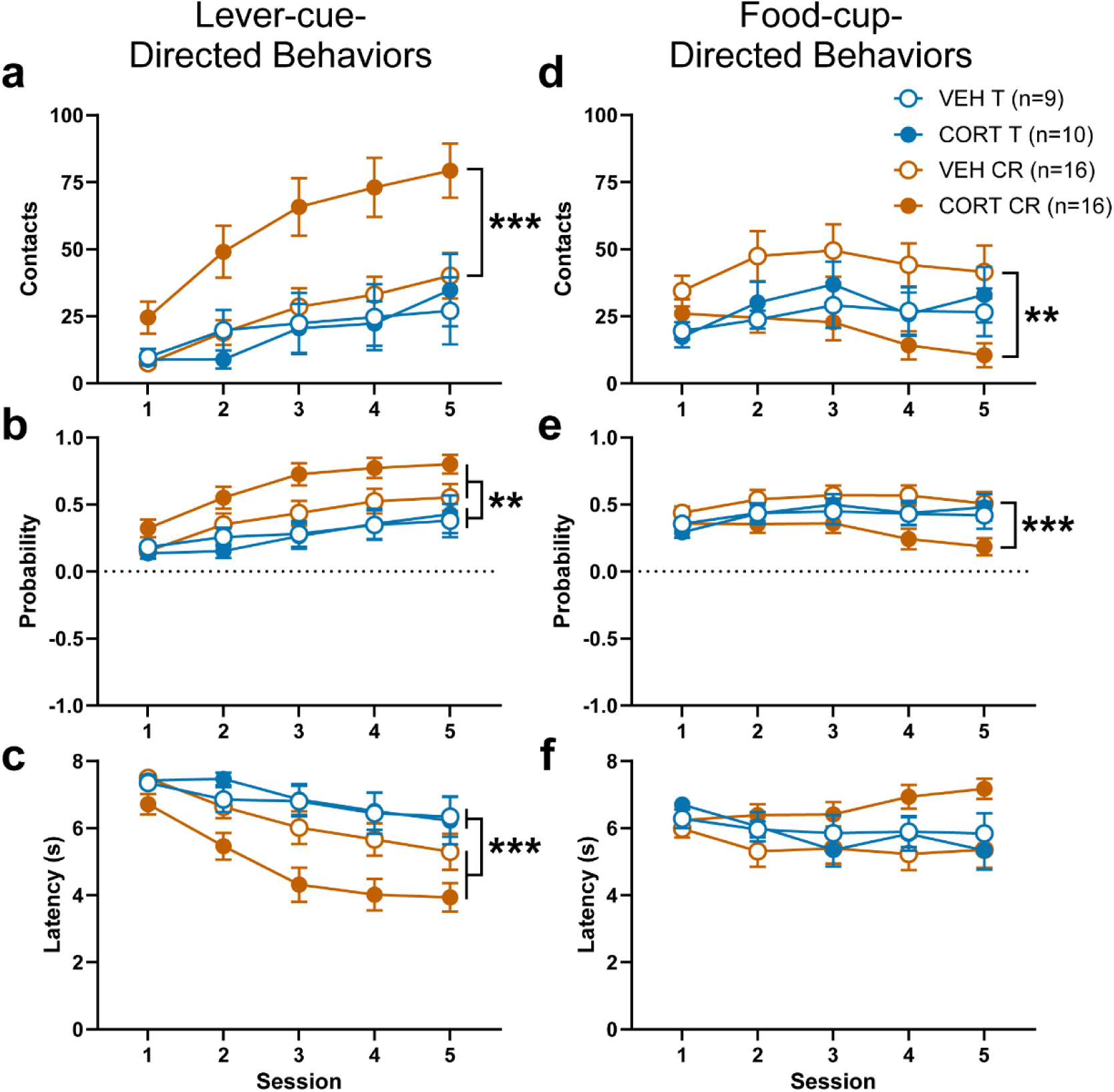
Male rats from Charles River exhibit more lever-cue-directed behaviors, and CORT treatment enhances the acquisition of these behaviors while attenuating food-cup-directed behaviors. Lever-cue-directed (a-c) and food-cup-directed (d-f) behaviors during PavCA sessions 1-5 for Experiment 1. Data are shown as mean ± SEM for (a,d) number of contacts, (b,e) probability, or (c,f) latency to approach the lever-cue (left panels) or food-cup (right panels). (b,c) Compared to male rats from Taconic, male rats from Charles River had a greater number of contacts with the lever-cue (effect of vendor: *p=0.006), greater probability to approach the lever-cue (effect of vendor: **p=0.003) and a decreased latency to approach the lever-cue (effect of vendor: ***p=0.002). Administration of CORT enhanced lever-cue contacts selectively in male rats from Charles River (treatment x vendor: p=0.031, *post hoc*: ***p=0.001). Similarly, relative to vehicle-treated rats, CORT administration selectively decreased food-cup contacts (treatment x vendor: p=0.033, *post hoc*: **p=0.003), and the probability to approach the food cup (treatment x vendor: p=0.035, *post hoc*: ***p=0.001) in male rats from Charles River.

**Figure 3.**
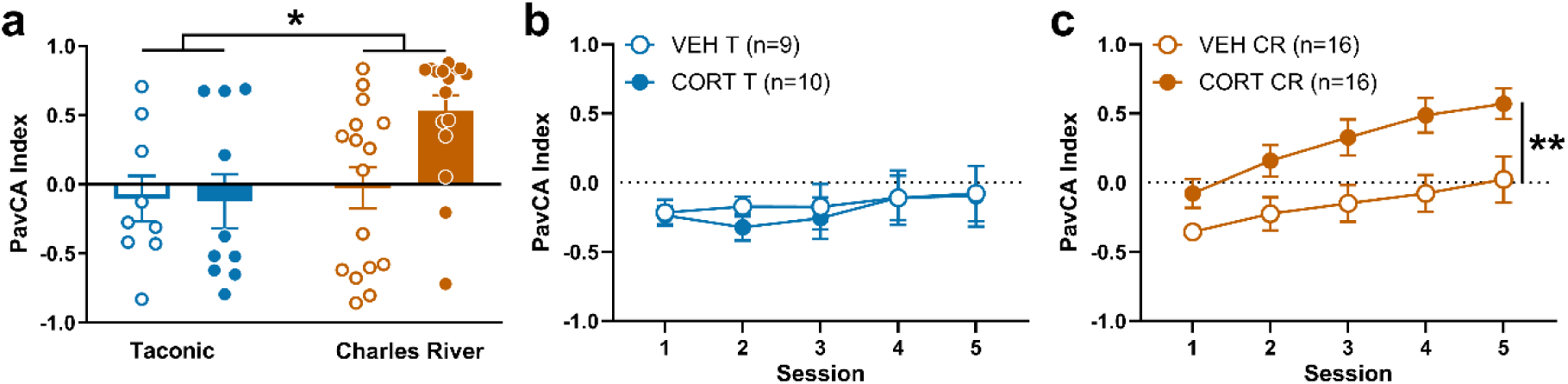
CORT enhances the propensity to sign-track selectively in male rats from Charles River. Effect of CORT or VEH administration on PavCA index in male rats from Taconic and Charles River in Experiment 1. (a) PavCA index (mean ± SEM) averaged across sessions 4 and 5. Charles River rats displayed greater sign-tracking behaviors than Taconic male rats (effect of vendor: *p=0.025). This effect appears to be primarily driven by increased sign-tracking in Charles River males treated with CORT (trend for vendor x treatment interaction: p=0.074). PavCA index (mean ± SEM) across sessions 1-5 in (b) male Taconic rats and (c) male Charles River rats. Administration of CORT increased the propensity to sign-track in (c) male rats from Charles River compared to their VEH counterparts (*post hoc*: **p=0.002) but CORT did not significantly affect the propensity to sign-track in (b) male rats from Taconic.

Behavior during the conditioned reinforcement test sessions in Experiments 1 and 2 (Figures 4 and 7) was assessed using an ANOVA with nose pokes, lever-cue contacts, and incentive value index as dependent variables. Analysis of nose poke responding included treatment and vendor as between-subjects factors and port as the within-subjects factor. Similar analyses were conducted for Experiment 3 (Figure 9) with the addition of sex as a between-subjects factor.

**Figure 4.**
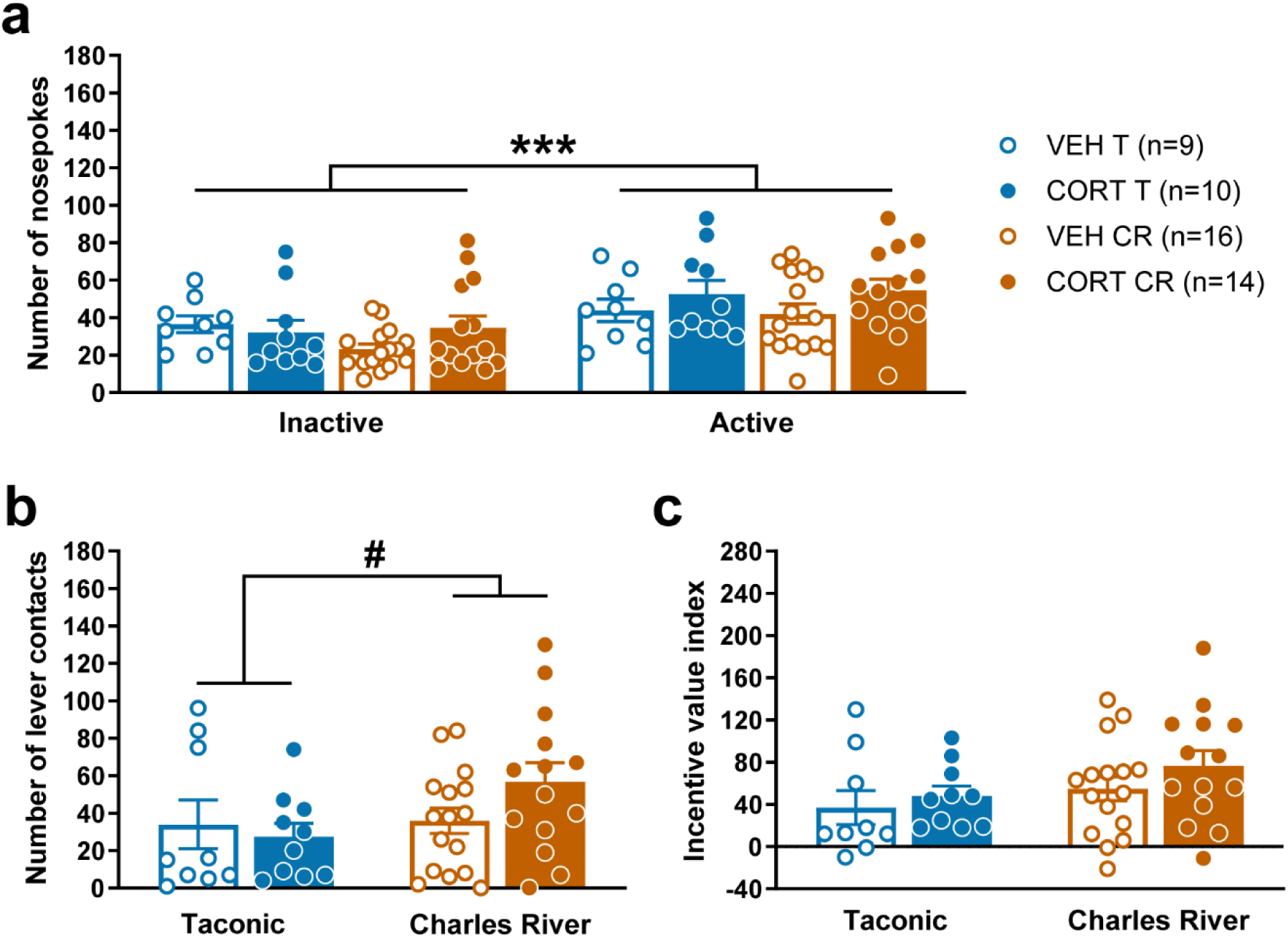
There are no significant effects of prior CORT treatment on the conditioned-reinforcing properties of the lever-cue for male rats. (a) Number of nose pokes (mean ± SEM) into the inactive and active ports for male rats from Taconic and Charles River in Experiment 1. All rats exhibited a greater number of nose pokes into the active port relative to the inactive port (effect of port: ***p<0.001). (b) Number of lever-cue contacts (mean ± SEM) for male rats from Taconic and Charles River. Compared to male rats from Taconic, male rats from Charles River had a tendency to exhibit a greater number of lever-cue contacts upon its presentation during the conditioned reinforcement test (trend for effect of vendor: #p=0.062). (c) Incentive value index (mean ± SEM) for male rats from Taconic and Charles River. There were no significant effects of vendor, treatment, or interaction on the incentive value index. Prior systemic CORT treatment did not significantly affect any of the measures during the conditioned reinforcement test.

## Results

### Experiment 1: Effect of CORT and vendor on the acquisition of PavCA in male rats

#### Pavlovian Conditioned Approach (PavCA)

The effects of CORT on lever-cue- and food-cup-directed behaviors across five PavCA sessions were analyzed for male rats from Charles River and Taconic. Table 2 includes the results of these analyses with key effects emphasized below. All of the rats showed a change in lever-cue-directed behaviors (Figure 2a-c; Table 2) across sessions (effect of session: lever-cue contacts F_4,47.760_=12.095, p<0.001; probability to approach the lever-cue F_4,84.699_=13.498, p<0.001; latency to approach the lever-cue F_4,47.687_=11.963, p<0.001), but the degree of sign-tracking behavior across sessions differed between rats from the two vendors (vendor x session: latency to approach the lever-cue F_4,47.687_=2.679, p=0.043). Relative to those from Taconic, male rats from Charles River exhibited a greater number of lever-cue contacts, an increased probability to approach the lever-cue, and a decreased latency to approach the lever-cue (Figure 2a-c; effect of vendor: lever-cue contacts F_1,47.092_=8.407, p=0.006; probability to approach the lever-cue F_1,47.477_=9.897, p=0.003; latency to approach the lever-cue F_1,47.067_=10.636, p=0.002). CORT administration enhanced these vendor effects. Specifically, as shown in Figure 2a, lever-cue-directed behavior increased in response to CORT selectively in male rats from Charles River (treatment x vendor: F_1,47.092_=4.934, p=0.031, *post hoc*: p=0.001).

**Table 2.**
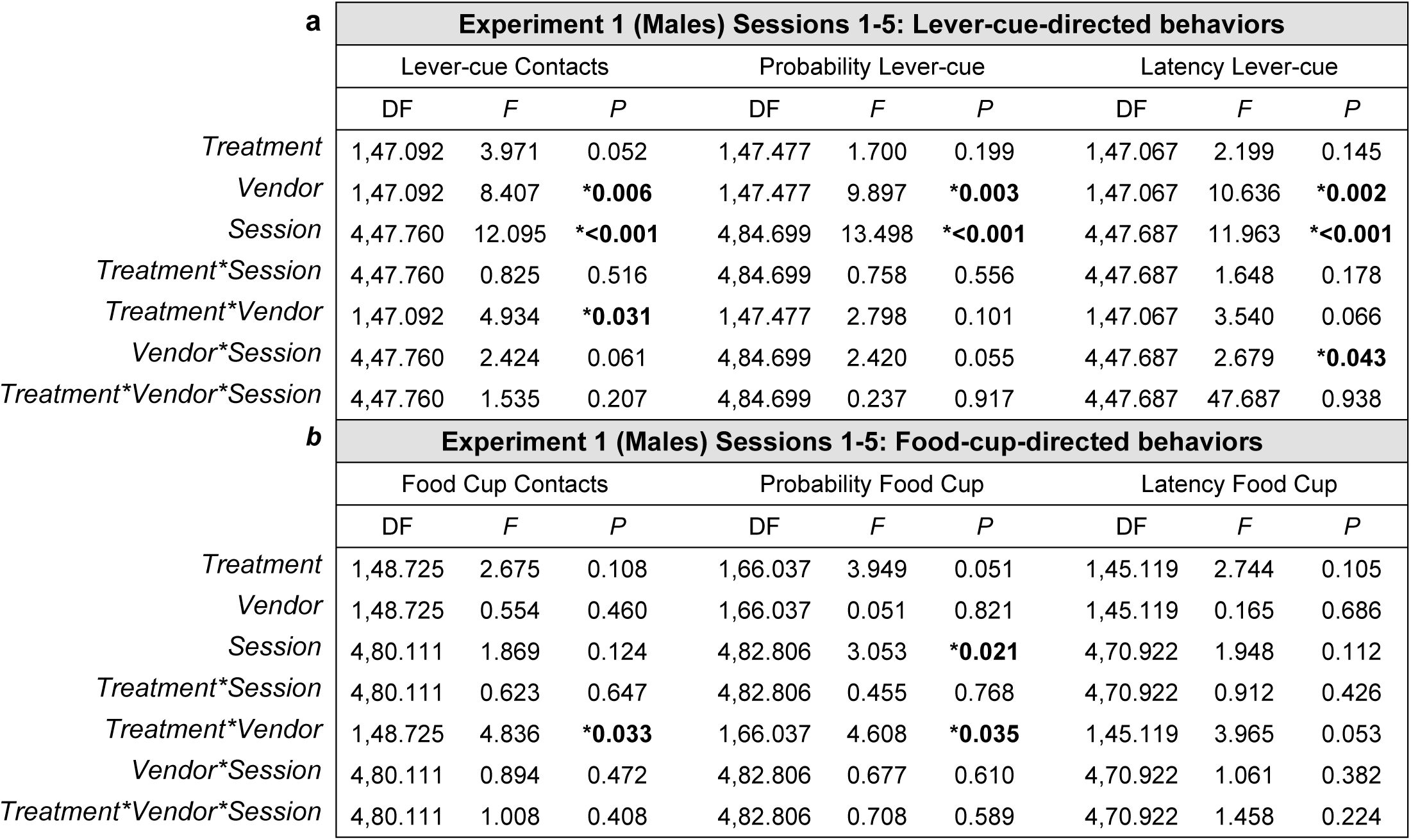
Statistical analyses for lever-cue- and food-cup-directed behaviors for Experiment 1. Data from linear mixed effects model analyses for sessions 1-5 for male rats in Experiment 1. Contacts, probability, and latency are represented for (a) lever-cue-directed behaviors and (b) food-cup-directed behaviors. Significant effects are indicated in bold with an asterisk.

In contrast to lever-cue-directed behaviors, there was not a significant effect of vendor for food-cup-directed behaviors in male rats (Figure 2d-f; Table 2), and only the probability to approach the food cup changed significantly across sessions (effect of session: probability to approach food cup F_4,82.806_=3.053, p=0.021; Figure 2e). There was, however, a significant interaction between treatment and vendor for the number of food cup contacts (F_1,48.725_=4.836, p=0.033; Figure 2d) and the probability to contact the food cup (F_1,66.037_=4.608, p=0.035; Figure 2e). *Post hoc* analyses revealed that CORT administration decreased the number of food cup contacts (F_1,48.581_=9.900, p=0.003; Figure 2d) and the probability to approach the food cup (F_1,65.757_=11.517, p=0.001; Figure 2e) in male rats from Charles River relative to VEH-treated controls from the same vendor. There was not a significant effect of CORT on goal-directed behavior in male rats from Taconic.

As shown in Figure 2 and described above, the administration of CORT enhanced the acquisition of sign-tracking behaviors, and attenuated the acquisition of goal-tracking behaviors, but did so selectively in male rats from Charles River. It is possible that these vendor-specific effects are due to inherent differences in the tendency to sign-track, as there was a significant effect of vendor on the average PavCA index across sessions 4 and 5 (F_1,47_=5.393, p=0.025; Figure 3a), with male rats from Charles River showing greater sign-tracking behavior. Although it did not reach statistical significance, there was a trend toward a vendor x treatment interaction (F_1,47_=3.340, p=0.074). Further analysis of PavCA index across sessions 1-5 revealed a significant effect of vendor (F_1,48.020_=4.877, p=0.032) and a significant vendor x treatment interaction (F_1,48.020_=4.973, p=0.030). To better illustrate this interaction, vendors are graphed separately in Figure 3b and 3c. In support of the data described above, CORT administration selectively increased the propensity to sign-track in male rats from Charles River (*post hoc:* F_1,47.81_=10.304, p=0.002, Figure 3c) and not those from Taconic (Figure 3b).

#### Conditioned Reinforcement Test

The conditioned reinforcing properties of the lever-cue were assessed in the same male rats from Taconic and Charles River. These rats received VEH or CORT prior to PavCA sessions1-5, but not immediately prior to the conditioned reinforcement test. All rats exhibited a greater number of nose pokes into the “active” port relative to the “inactive” port (effect of port: F_1,45_=30.084, p<0.001; Figure 4a). There were no significant effects of treatment, vendor, nor an interaction for nose-poke responding, lever presses, or the incentive value index (Figure 4). However, male rats from Charles River tended to exhibit a greater number of lever-cue contacts upon its presentation during the conditioned reinforcement test compared to male rats from Taconic (trending effect of vendor: F_1,45_=3.653, p=0.062; Figure 4B). While not statistically significant, this trend is consistent with the vendor effects on sign-tracking behavior described above.

### Experiment 2: Effect of CORT and vendor on the acquisition of PavCA in female rats

#### Pavlovian Conditioned Approach (PavCA)

The effect of CORT on lever-cue- and food-cup-directed behaviors across five PavCA sessions were analyzed for female rats from Charles River (n=28) and Taconic (n=30). Table 3 includes the results of these analyses with key effects emphasized below.

**Table 3.** Statistical analyses for lever-cue- and food-cup-directed behaviors for Experiment 2. Data from linear mixed effects model analyses for sessions 1-5 for female rats in Experiment 2. Contacts, probability, and latency are represented for (a) lever-cue-directed behaviors and (b) food-cup-directed behaviors. Significant effects are indicated in bold with an asterisk.

| *a* | **Experiment 2 (Females) Sessions 1-5: Lever-cue-directed behaviors** |  |  |  |  |  |  |  |  |
| --- | --- | --- | --- | --- | --- | --- | --- | --- | --- |
|  | Lever-cue Contacts |  |  | Probability Lever-cue |  |  | Latency Lever-cue |  |  |
|  | DF | *F* | *P* | DF | *F* | *P* | DF | *F* | *P* |
| *Treatment* | 1,54.995 | 1.983 | 0.165 | 1,54.950 | 2.893 | 0.095 | 1,53.314 | 3.670 | 0.061 |
| *Vendor* | 1,54.995 | 9.416 | **\*0.003** | 1,54.950 | 9.040 | **\*0.004** | 1,53.314 | 11.806 | **\*0.001** |
| *Session* | 4,86.035 | 22.076 | **\*<0.001** | 4,92.728 | 30.701 | **\*<0.001** | 4,76.183 | 30.769 | **\*<0.001** |
| *Treatment\*Session* | 4,86.035 | 1.305 | 0.275 | 4,92.728 | 1.146 | 0.340 | 4,76.183 | 1.682 | 0.163 |
| *Treatment\*Vendor* | 1,54.995 | 0.036 | 0.850 | 1,54.950 | 0.221 | 0.640 | 1,53.314 | 0.385 | 0.538 |
| *Vendor\*Session* | 4,86.035 | 4.587 | **\*0.002** | 4,92.728 | 5.149 | **\*0.001** | 4,76.183 | 5.946 | **\*<0.001** |
| *Treatment\*Vendor\*Session* | 4,86.035 | 0.356 | 0.839 | 4,92.728 | 0.278 | 0.892 | 4,76.183 | 0.669 | 0.616 |
| *b* | **Experiment 2 (Females) Sessions 1-5: Food-cup-directed behaviors** |  |  |  |  |  |  |  |  |
|  | Food Cup Contacts |  |  | Probability Food Cup |  |  | Latency Food Cup |  |  |
|  | DF | *F* | *P* | DF | *F* | *P* | DF | *F* | *P* |
| *Treatment* | 1,55.761 | 3.599 | 0.063 | 1,55.713 | 3.263 | 0.076 | 1,55.736 | 4.351 | **\*0.042** |
| *Vendor* | 1,55.761 | 2.744 | 0.103 | 1,55.713 | 4.566 | **\*0.037** | 1,55.736 | 4.242 | **\*0.044** |
| *Session* | 4,89.454 | 4.221 | **\*0.004** | 4,95.648 | 5.047 | **\*<0.001** | 4,89.984 | 4.954 | **\*0.001** |
| *Treatment\*Session* | 4,89.454 | 2.212 | 0.074 | 4,95.648 | 1.963 | 0.106 | 4,89.984 | 1.809 | 0.134 |
| *Treatment\*Vendor* | 1,55.761 | 0.007 | 0.934 | 1,55.713 | 0.003 | 0.954 | 1,55.736 | 0.008 | 0.930 |
| *Vendor\*Session* | 4,89.454 | 3.400 | **\*0.012** | 4,95.648 | 6.053 | **\*<0.001** | 4,89.984 | 4.226 | **\*0.004** |
| *Treatment\*Vendor\*Session* | 4,89.454 | 0.160 | 0.958 | 4,95.648 | 0.314 | 0.868 | 4,89.984 | 0.435 | 0.783 |

Similar to the male rats, female rats exhibited a change in lever-directed behaviors across sessions (effect of session: lever-cue contacts F_4,86.035_=22.076, p<0.001; probability to approach the lever-cue F_4,92.728_=30.701, p<0.001; latency to approach the lever-cue F_4,76.183_=30.769, p<0.001) with the degree of sign-tracking behavior varying across sessions between the two vendors (vendor x session: lever-cue contacts F_4,86.035_=4.587, p=0.002; probability to approach the lever-cue F_4,92.728_=5.149, p=0.001; latency to approach the lever-cue F_4,76.183_=5.946, p<0.001). As shown in Figure 5, relative to those from Taconic, female rats from Charles River exhibited a greater number of lever-cue contacts, an increased probability to approach the lever-cue, and a greater latency to approach the lever-cue (Figure 5a-c; effect of vendor: lever-cue contacts F_1,54.995_=9.416, p=0.003; probability to approach the lever-cue F_1,54.950_=9.040, p=0.004; latency to approach the lever-cue F_1,53.314_=11.806, p=0.001). There were no significant effects of treatment on sign-tracking behaviors for female rats (see Table 3); however, females treated with CORT tended to have an increased probability (trending effect of treatment: F_1,54.950_=2.893, p=0.095; Figure 5b) and decreased latency to approach the lever-cue (trending effect of treatment: F_1,53.314_=3.670, p=0.061; Figure 5c). While not statistically significant, this trend is consistent with the effects of CORT treatment seen in males from Charles River as described in Experiment 1.

**Figure 5.**
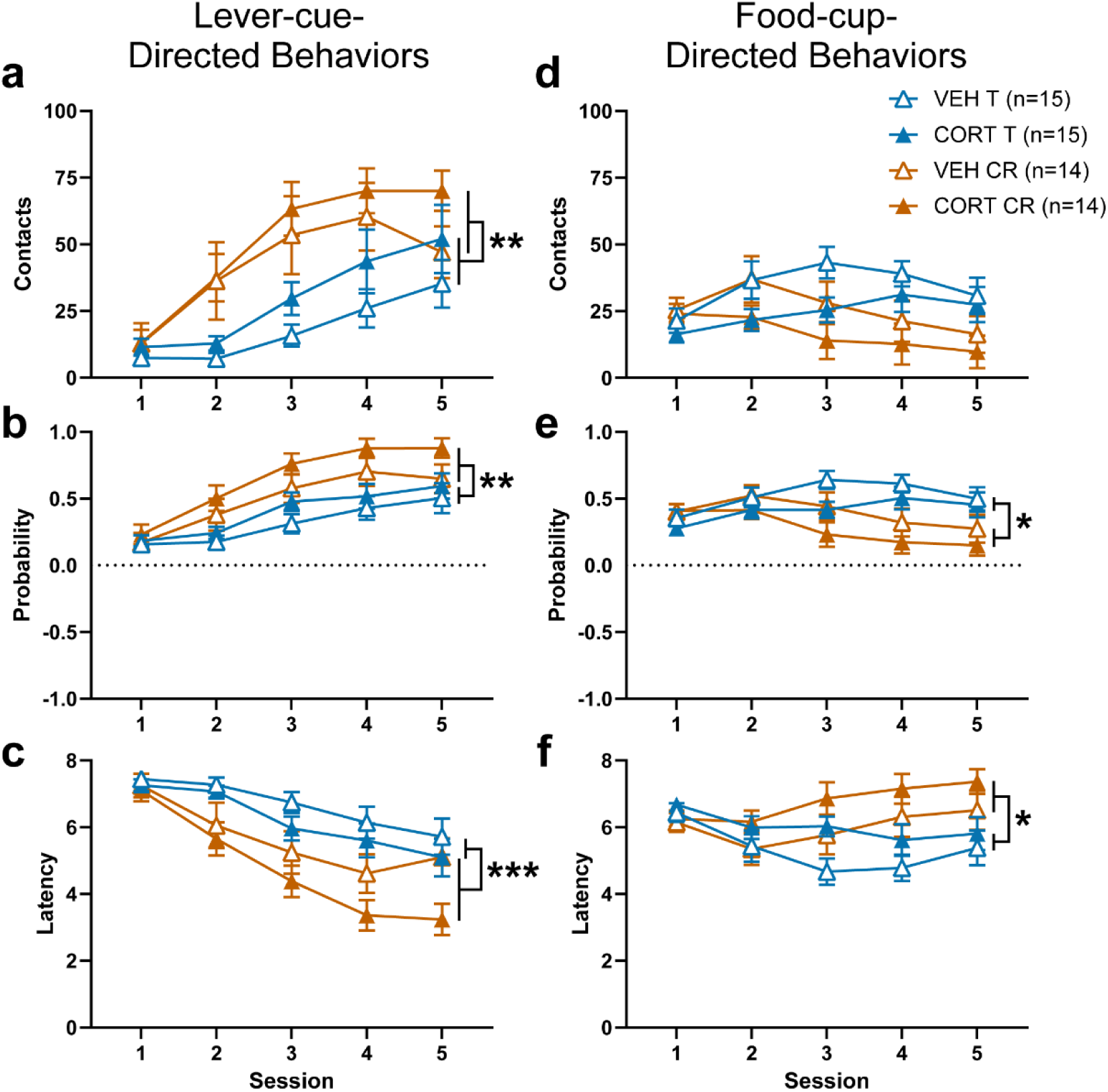
Female rats from Charles River exhibit more lever-cue-directed and less food-cup-directed behavior during acquisition of PavCA relative to female rats from Taconic. Lever-cue-directed (a-c) and food-cup-directed (d-f) behaviors during PavCA sessions 1-5 for Experiment 2. Data are shown as mean ± SEM for (a,d) number of contacts, (b,e) probability, or (c,f) latency to approach the lever-cue (left panels) or food cup (right panels). (a-c) Female rats from Charles River exhibited a greater number of lever-cue contacts (effect of vendor: **p=0.003), an increased probability to approach the lever-cue (effect of vendor: **p=0.004), and an increased latency to approach the lever-cue (effect of vendor: ***p=0.001) compared to female rats from Taconic. (d-f) Compared to female rats from Taconic, female rats from Charles River exhibited a decreased probability to approach the magazine (effect of vendor: *p=0.037) and an increased latency to approach the magazine (effect of vendor: *p=0.044). Regardless of vendor, administration of CORT in female rats significantly increased the latency to approach the food cup (effect of treatment: p=0.042, significance not shown) and had a tendency to attenuate other food-cup-directed behaviors, while increasing lever-cue-directed behaviors (trending effects not shown).

While food-cup-directed behavior changed across sessions for all rats (Figure 5d-f; effect of session: food-cup contacts F_4,89.454_=4.221, p=0.004; probability to approach the food cup F_4,95.648_=5.047, p<0.001; latency to approach the food cup F_4,89.984_, p=0.001), the degree of goal-tracking behavior varied across sessions by vendor (vendor x session: food cup contacts F_4,89.454_=3.400, p=0.012; probability to approach the food cup F_4,95.648_=6.053, p<0.001; latency to approach the food cup F_4,89.984_=4.226, p=0.004). Female rats from Charles River exhibited less goal-tracking behavior relative to those from Taconic (Figure 5e,f; effect of vendor: probability to approach the food cup: F_1,55.713_=4.566, p=0.037; latency to approach the food cup F_1,55.736_=4.242, p=0.044). Administration of CORT significantly increased the latency to approach the food cup (effect of treatment: F_1,55.736_=4.351, p=0.042; Figure 5f) and tended to decrease the number of food cup contacts (trending effect of treatment: F_1,55.761_=3.599, p=0.063; Figure 5d) and probability to approach the food cup (trending effect of treatment: F_1,55.713_=3.263, p=0.076; Figure 5e). Thus, similar to the effects reported for male rats from Charles River in Experiment 1, CORT treatment tended to attenuate goal-tracking behavior in female rats, although the effect of CORT in females did not depend on vendor.

In agreement with the data described above, analysis of the average PavCA index across sessions 4-5 revealed a significant effect of vendor (effect of vendor: F_1,54_=11.264, p=0.001), but no significant interaction of vendor and treatment. Female rats from Charles River exhibited a greater tendency to sign-track, regardless of treatment (Figure 6a). When the PavCA index was analyzed across sessions 1-5, female rats from Charles River had a greater tendency to sign-track than those from Taconic (effect of vendor: F_1,55.050_=8.402, p=0.005). In contrast to male rats, the effect of CORT did not depend on vendor in female rats as there was not a significant interaction between vendor and treatment; rather, CORT administration increased the tendency to sign-track in female rats, regardless of vendor (effect of treatment: F_1,55.050_=4.620, p=0.036; Figure 6b,c).

**Figure 6.**
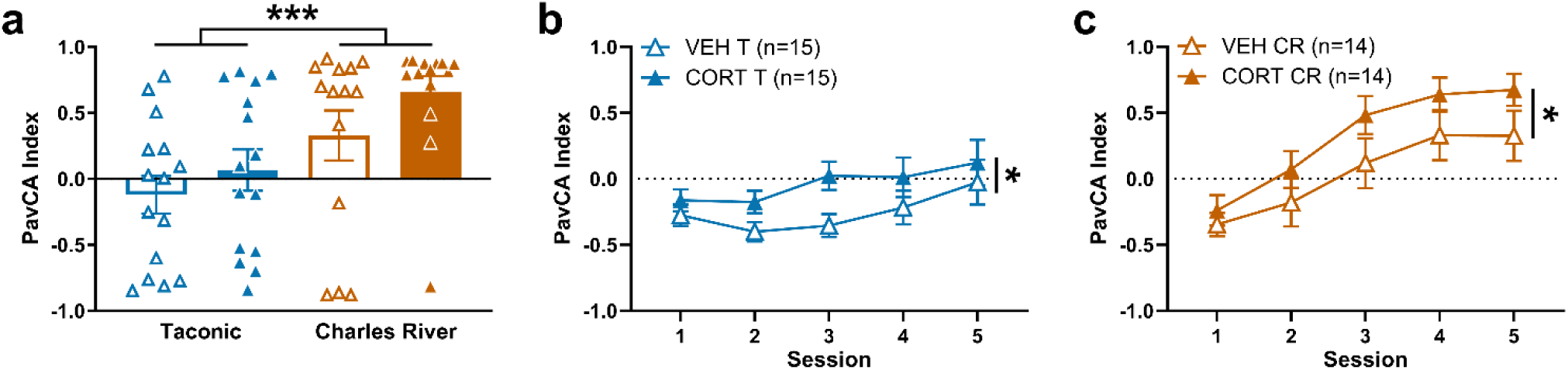
Relative to female rats from Taconic, those from Charles River have a greater propensity to sign-track, and CORT enhances the propensity to sign-track in female rats across vendors. Effect of CORT or VEH administration on PavCA index in female rats from Taconic and Charles River in Experiment 2. (a) PavCA index (mean ± SEM) averaged across sessions 4 and 5. Charles River female rats displayed greater sign-tracking behavior than Taconic female rats, regardless of treatment (effect of vendor: ***p=0.001). PavCA index (mean ± SEM) across sessions 1-5 in (b) female Taconic rats and (c) female Charles River rats. Administration of CORT increased the propensity to sign-track in female rats compared to those treated with VEH, regardless of vendor (effect of treatment: *p=0.036). There were no significant vendor x treatment interactions for any of these measures.

#### Conditioned Reinforcement Test

The conditioned reinforcing properties of the lever-cue were assessed in the same female rats from Taconic and Charles River (Figure 7). These rats received VEH or CORT prior to PavCA sessions 1-5, but not immediately prior to the conditioned reinforcement test. All rats exhibited a greater number of nose pokes into the “active” port relative to the “inactive” port (effect of port: F_1,54_=30.867, p<0.001; Figure 7a). There were no significant effects of treatment, vendor, nor an interaction on nose poke responding. There were no significant effects of treatment nor interactions between treatment and vendor for lever-cue contacts or the incentive value index. However, female rats from Charles River exhibited a greater number of lever-cue contacts upon its presentation during the conditioned reinforcement test (effect of vendor: F_1,54_=6.546, p=0.013; Figure 7b), and had a greater incentive value index compared to Taconic female rats (effect of vendor: F_1,54_=7.096, p=0.010; Figure 7c). This vendor effect is consistent with the vendor effects on sign-tracking described above.

**Figure 7.**
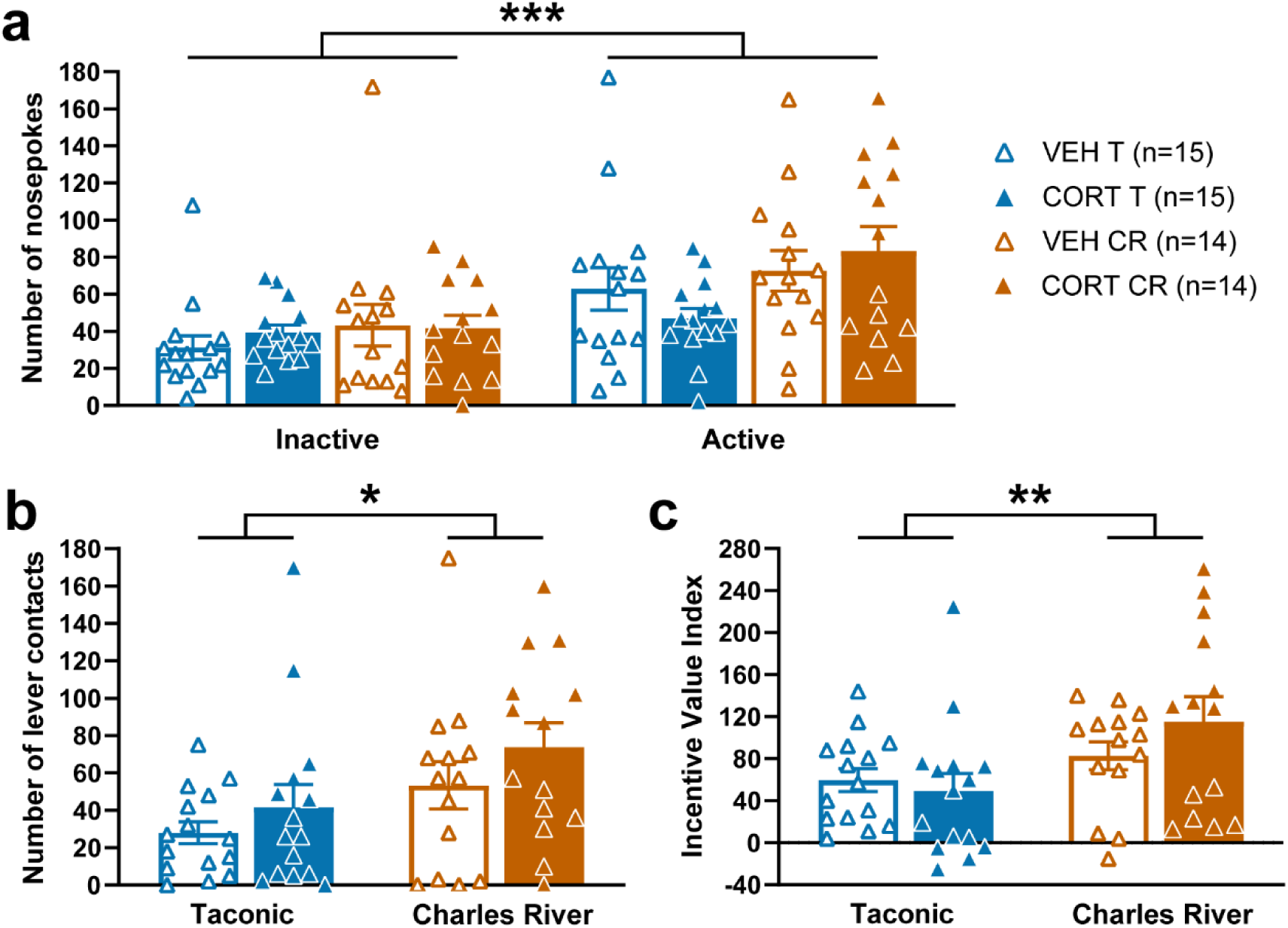
The conditioned-reinforcing properties of the lever-cue are enhanced in female rats from Charles River compared to those from Taconic. (a) Number of nose pokes (mean ± SEM) into the inactive and active ports for female rats from Taconic and Charles River in Experiment 2. All rats exhibited a greater number of nose pokes into the active port relative to the inactive port (effect of port: ***p<0.001). (b) Number of lever-cue contacts (mean ± SEM) for female rats from Taconic and Charles River. Female rats from Charles River exhibited a greater number of lever-cue contacts upon its presentation during the conditioned reinforcement test compared to those from Taconic (effect of vendor: *p=0.013). (c) Incentive value index (mean ± SEM) for female rats from Taconic and Charles River. Charles River female rats had a greater incentive value index compared to Taconic female rats (effect of vendor: **p=0.010). Prior systemic CORT treatment did not significantly affect any of the measures during the conditioned reinforcement test.

### Experiment 3: Effect of CORT, vendor, and sex on the expression of PavCA

#### Pavlovian Conditioned Approach (PavCA)

In Experiment 3, males and females were run concurrently and CORT treatment did not begin until session 6, allowing the assessment of the influence of sex and vendor on inherent tendencies in acquisition of conditioned responding in the absence of treatment (i.e., sessions 1-5; Figure 8). Analyses of the effects of CORT on the expression of conditioned responding were performed on sessions 6-9 (Figure 8). Table 4 includes the results of these analyses with key effects emphasized below.

**Figure 8.**
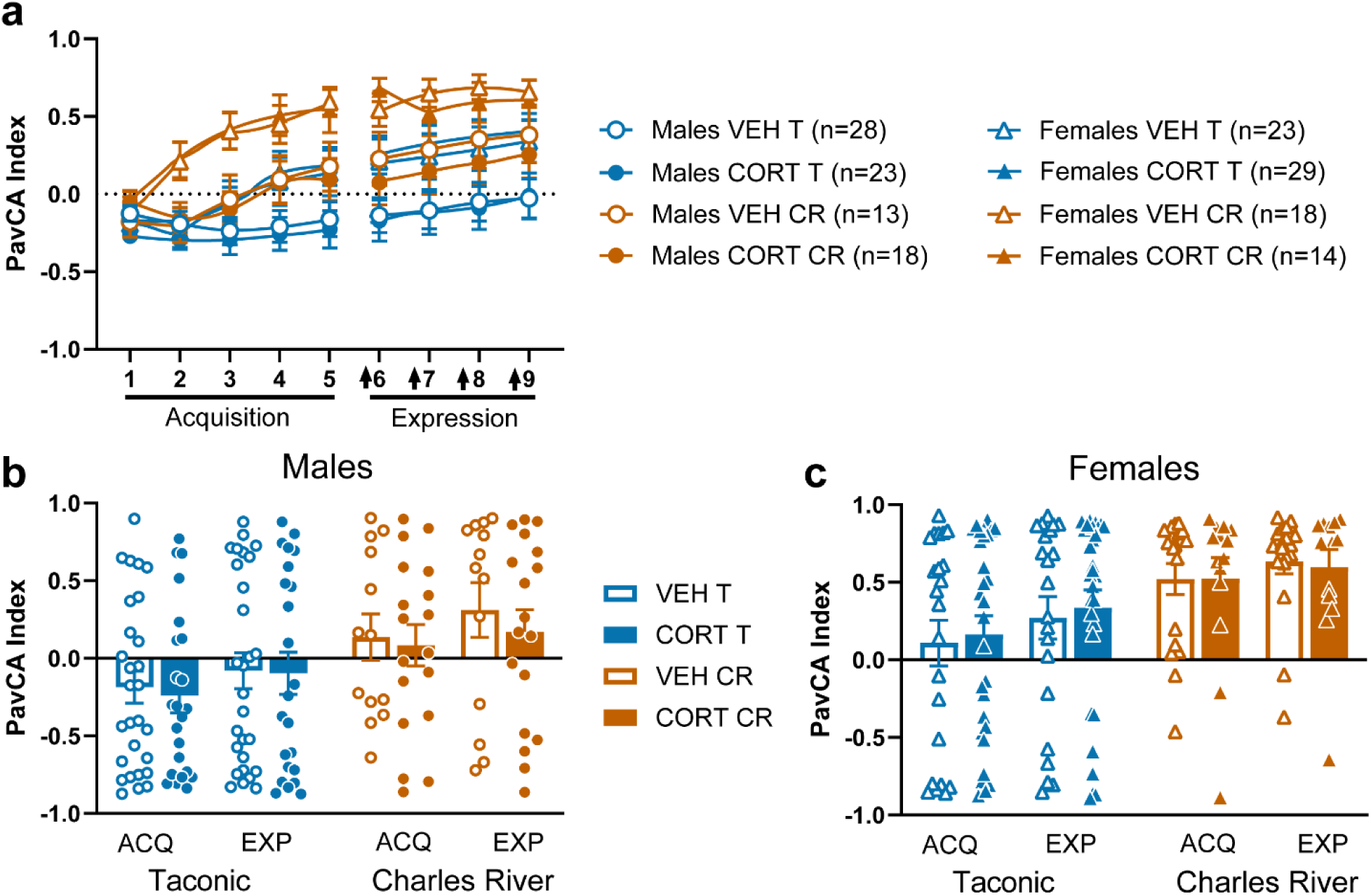
CORT did not affect the expression of a conditioned response in male or female rats from Charles River or Taconic. (a) PavCA Index (mean ± SEM) across sessions 1-5 (acquisition) and sessions 6-9 (expression) in Experiment 3. Rats from Charles River displayed greater sign-tracking behaviors compared to rats from Taconic both before (sessions 1-5; effect of vendor: p<0.001) and after treatment (sessions 6-9; effect of vendor: p=0.001). Female rats exhibited greater sign-tracking than male rats before treatment (sessions 1-5; effect of sex: p<0.001) and this sex difference persisted during treatment (sessions 6-9; effect of sex: p<0.001). Administration of systemic CORT on sessions 6-9 (indicated by the black arrows) did not significantly impact PavCA Index. PavCA Index (mean ± SEM) averaged across sessions 4 and 5 for the acquisition (ACQ) phase and averaged across sessions 6-9 for the expression (EXP) phase for (b) male and (c) female rats in Experiment 3. PavCA indices were greater during the expression phase than the acquisition phase (effect of phase: p<0.001). Regardless of treatment or vendor, female rats displayed greater sign-tracking behavior compared to male rats across both phases (effect of sex: p<0.001). Additionally, rats from Charles River displayed greater sign-tracking behavior compared to rats from Taconic across both phases (effect of vendor: p<0.001). There were no significant interactions of vendor, sex, or phase and no main effect or interactions with treatment.

**Table 4.** Statistical analyses for PavCA index across sessions during acquisition and expression of PavCA for Experiment 3. Data from linear mixed effects model analyses for PavCA index across (a) sessions 1-5 and (b) sessions 6-9 for rats of both sexes in Experiment 3. Rats did not receive treatment until session 6-9, so the treatment factor in session 1-5 refers to eventual treatment group. Significant effects are indicated in bold with an asterisk.

| **a** | **Experiment 3 Acquisition**<br>**Sessions 1-5** |  |  |
| --- | --- | --- | --- |
|  | PavCA Index |  |  |
|  | DF | F | P |
| *Treatment* | 1,160.181 | 0.023 | 0.879 |
| *Vendor* | 1,160.181 | 16.162 | **\*<0.001** |
| *Sex* | 1,160.181 | 16.191 | **\*<0.001** |
| *Session* | 4,286.416 | 14.176 | **\*<0.001** |
| *Treatment\*Vendor* | 1,160.181 | 0.079 | 0.779 |
| *Treatment\*Sex* | 1,160.181 | 0.214 | 0.644 |
| *Treatment\*Session* | 4,284.416 | 0.991 | 0.413 |
| *Vendor\*Sex* | 1,160.181 | 1.194 | 0.276 |
| *Vendor\*Session* | 4,286.416 | 2.713 | **\*0.030** |
| *Sex\*Session* | 4,286.416 | 5.377 | **\*<0.001** |
| *Treatment\*Vendor\*Sex* | 1,160.181 | 0.108 | 0.743 |
| *Treatment\*Vendor\*Session* | 4,286.416 | 0.393 | 0.814 |
| *Treatment\*Sex\*Session* | 4,286.416 | 0.149 | 0.963 |
| *Vendor\*Sex\*Session* | 4,286.416 | 3.237 | **\*0.013** |
| *Treatment\*Vendor\*Sex\*Session* | 4,286.416 | 0.489 | 0.744 |
| **b** | **Experiment 3 Expression**<br>**Sessions 6-9** |  |  |
|  | PavCA Index |  |  |
|  | DF | F | P |
| *Treatment* | 1,158.159 | 0.148 | 0.701 |
| *Vendor* | 1,158.159 | 11.194 | **\*0.001** |
| *Sex* | 1,158.159 | 16.756 | **\*<0.001** |
| *Session* | 3,317.957 | 5.768 | **\*<0.001** |
| *Treatment\*Vendor* | 1,158.159 | 0.307 | 0.580 |
| *Treatment\*Sex* | 1,158.159 | 0.202 | 0.654 |
| *Treatment\*Session* | 3,317.957 | 0.724 | 0.539 |
| *Vendor\*Sex* | 1,158.159 | 0.018 | 0.894 |
| *Vendor\*Session* | 3,317.957 | 0.615 | 0.606 |
| *Sex\*Session* | 3,317.957 | 0.708 | 0.548 |
| *Treatment\*Vendor\*Sex* | 1,158.159 | 0.011 | 0.918 |
| *Treatment\*Vendor\*Session* | 3,317.957 | 1.498 | 0.215 |
| *Treatment\*Sex\*Session* | 3,317.957 | 0.902 | 0.440 |
| *Vendor\*Sex\*Session* | 3,317.957 | 0.708 | 0.548 |
| *Treatment\*Vendor\*Sex\*Session* | 3,317.957 | 1.434 | 0.233 |

Linear mixed model analysis of the PavCA index during acquisition across sessions 1-5 (Figure 8a) revealed no significant main effects or interactions with treatment, indicating that the counterbalancing of treatment groups based on PavCA index prior to the beginning of treatment was successful. Rats from Charles River displayed greater sign-tracking behaviors compared to rats from Taconic (effect of vendor: F_1,160.181_=16.162, p<0.001). This vendor difference varied by sex and session (vendor x sex x session: F_4,286.416_=3.237, p=0.013) with the vendor difference arising earlier in females (session 2-5, *post hoc*: p<0.001, p<0.001, p=0.007, and p=0.003, respectively) than in males (by session 4-5, *post hoc*: p=0.019 & p=0.016, respectively). Females exhibited greater sign-tracking behavior than males (effect of sex: F_1,160.181_=16.191, p<0.001). This sex difference was present in rats from both vendors but emerged slightly earlier in rats from Charles River (session 2-5, *post hoc*: p<0.001, p<0.001, p=0.009, and p=0.005, respectively) than rats from Taconic (session 3-5, *post hoc*: p=0.032, p=0.003, and p=0.002, respectively). These findings further demonstrate that rats from Charles River and female rats have a greater propensity to sign-track than rats from Taconic and male rats, respectively, with females from Charles River displaying the fastest and greatest acquisition of sign-tracking (Figure 8a).

The effects of CORT on the expression of a conditioned response in male and female rats from Charles River and Taconic were compared by analyzing PavCA index across sessions 6-9 (Figure 8a). Administration of systemic CORT did not significantly impact expression of sign-tracking behaviors as there were no significant main effects or interactions with treatment. There were significant main effects of vendor (F_1,158.159_=11.194, p=0.001) and sex (F_1,158.159_=16.756, p<0.001), but no significant interaction of vendor and sex, reflecting that rats from Charles River and female rats express more sign-tracking behavior than rats from Taconic and male rats, respectively. These sex and vendor differences are the same as described above during the acquisition sessions and likely reflect differences in inherent tendencies towards sign-tracking.

A repeated measures ANOVA was conducted on the average PavCA index to compare sign-tracking behavior *acquired* by PavCA sessions 4 and 5 with the sign-tracking behavior that was *expressed* in PavCA sessions 6-9 when treatment was given (Figure 8b and 8c). There were no significant effects of treatment and no significant interactions between any of the factors. Rats displayed significantly greater sign-tracking behavior during the expression phase compared to the acquisition phase of the experiment (effect of phase: F_1,158_=28.219, p<0.001). There was also a significant main effect of sex (effect of sex: F_1,158_=11.435, p<0.001), in which females expressed greater sign-tracking behavior compared to males across both phases; and a significant main effect of vendor (effect of vendor: F_1,158_=13.786, p<0.001), in which rats from Charles River expressed greater sign-tracking behavior compared to those from Taconic across both phases. This vendor effect is consistent with those reported above for Experiments 1 and 2, indicating that rats from Charles River tend to be skewed more towards sign-trackers relative to those from Taconic.

#### Conditioned Reinforcement Test

The conditioned reinforcing properties of the lever-cue were assessed in the same rats of both sexes from Taconic and Charles River (Figure 9). These rats received VEH or CORT prior to PavCA sessions 6-9, but not immediately prior to the conditioned reinforcement test. All rats exhibited a greater number of nose pokes into the “active” port relative to the “inactive” port (effect of port: F_1,152_=201.504, p<0.001; Figure 9a,d). A significant port by vendor interaction (F_1,152_=16.912, p<0.001) indicated that rats from Charles River had a greater number of nose pokes in the “active” port relative to those from Taconic (*post hoc*: p<0.001). There was also a port by sex interaction (F_1,152_=24.753, p<0.001), revealing that females poked more in the “active” port than males (*post hoc*: p<0.001). Regardless of sex, rats from Charles River exhibited a greater number of lever-cue contacts upon its presentation during the conditioned reinforcement test compared to rats from Taconic (effect of vendor: F_1,152_=43.739, p<0.001; Figure 9b,e), and females contacted the lever-cue more than males (effect of sex: F_1,152_=18.523, p<0.001). Rats from Charles River and females had greater incentive value indices relative to those from Taconic and males, respectively (effect of vendor: F_1,152_=39.354, p<0.001; effect of sex: F_1,152_=26.068, p<0.001; Figure 9c,f). These vendor and sex effects are consistent with those described above and the lack of significant effects (or interactions) of *prior* CORT treatment on any of the conditioned reinforcement test measures is consistent with Experiments 1 and 2.

**Figure 9.**
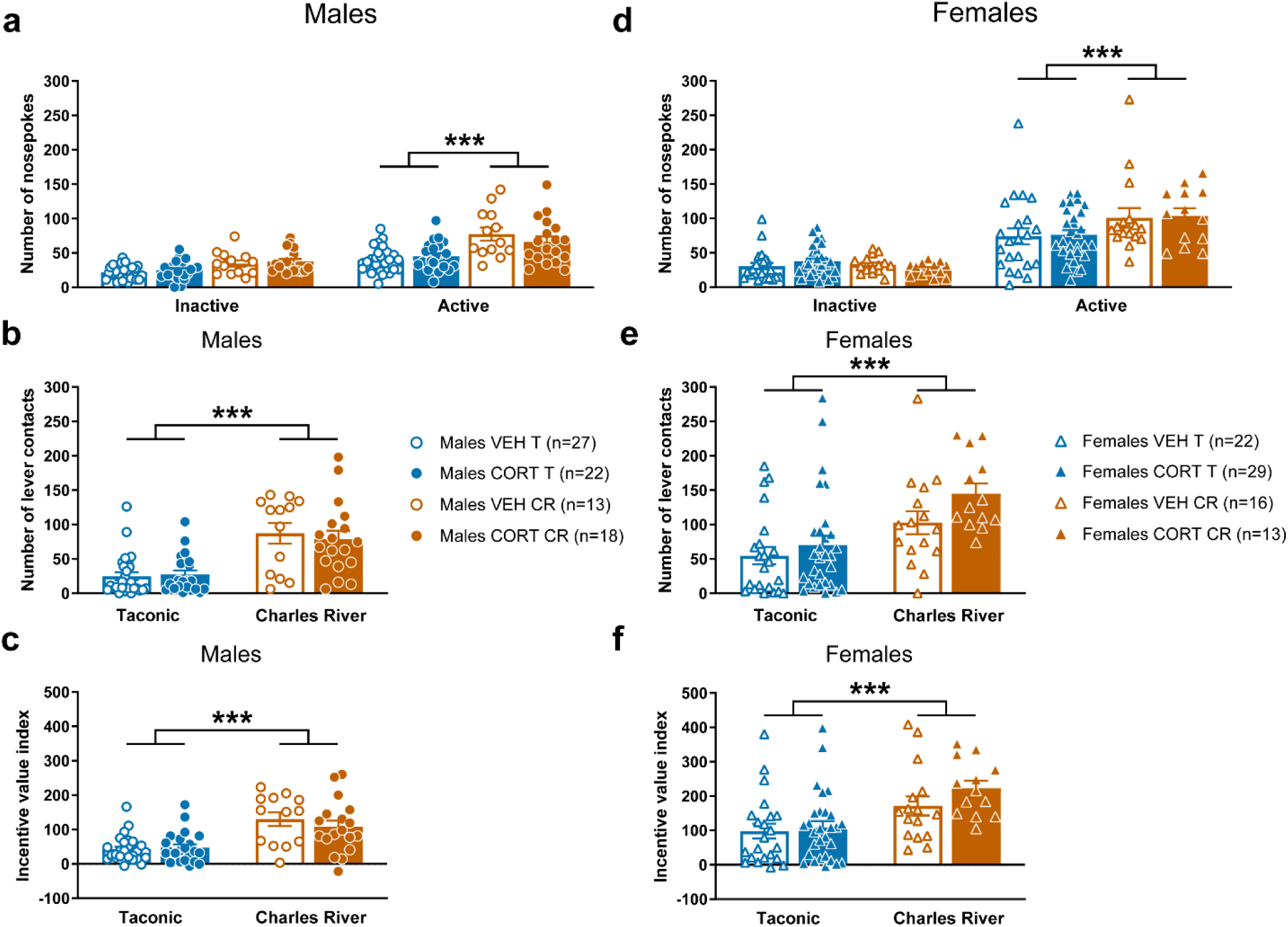
The conditioned-reinforcing properties of the lever-cue are enhanced in females compared to males, and in rats from Charles River compared to those from Taconic. Number of nose pokes (mean ± SEM) into the inactive and active ports for (a) male and (d) female rats from Taconic and Charles River in Experiment 3. Female rats poked more in the active port than males (*post hoc*: p<0.001), and rats from Charles River (orange) poked more in active port than those from Taconic (blue) (*post hoc*: ***p<0.001). All rats exhibited a greater number of nose pokes into the active port relative to the inactive port (effect of port: p<0.001). Number of lever-cue contacts (mean ± SEM) for (b) male and (e) female rats from Taconic and Charles River. Females exhibited a greater number of lever-cue contacts upon its presentation during the conditioned reinforcement test compared to males (effect of sex: p<0.001). Rats from Charles River contacted the lever-cue more than those from Taconic (effect of vendor: ***p<0.001). Incentive value index (mean ± SEM) for (c) male and (f) female rats from Taconic and Charles River. Charles River rats had a greater incentive value index compared to Taconic rats (effect of vendor: ***p<0.001), and females had a greater incentive value index than males (effect of sex: p<0.001). Prior systemic CORT treatment did not significantly affect any of the measures during the conditioned reinforcement test. There were no significant interactions with vendor or sex on any of the measures.

## Discussion

The current study revealed that systemic administration of corticosterone (CORT) influenced the propensity to attribute incentive value to reward cues, and that it did so in a sex- and vendor-dependent manner. Our overall hypothesis that CORT administration during training would increase sign-tracking behavior was only partially supported. CORT administration did influence the acquisition of sign-tracking behavior in male and female rats, but the effect in males was dependent on vendor. In the experiments conducted with males, CORT elicited an increase in the acquisition of sign-tracking behavior in rats from Charles River, but not in those from Taconic. Similarly, in experiments conducted with females, CORT elicited an increase in the acquisition of sign-tracking behavior, but the effect in females was not vendor-dependent. There were no significant effects of CORT treatment on the expression of sign-tracking behavior, but the effects of sex and vendor observed in the absence of treatment were consistent between all three experiments, suggesting that rats from Charles River have a greater tendency to sign-track relative to those from Taconic. Further, consistent with prior reports, female rats tended to sign-track more than male rats (Hughson et al., 2019; Pitcher et al., 2015).

Our hypothesis that administration of CORT would increase the propensity to sign-track in rats was derived largely from the results of studies with quail. Rice and colleagues found that administration of a glucocorticoid receptor (GR) antagonist (PT150) attenuated sign-tracking behaviors in male Japanese quail (Rice et al., 2019). Considering a GR antagonist attenuated sign-tracking, we expected administration of CORT, a GR agonist, to promote sign-tracking. Further, we have previously shown that, following a single PavCA session, peripheral CORT increases to a greater extent in male sign-trackers than male goal-trackers (Flagel et al., 2009). With these previous findings in mind, we used a pharmacological approach to investigate the role of CORT in incentive vs. predictive learning in rats. Our results did not fully support our hypothesis, as we found that administration of CORT did not increase the propensity to sign-track in all rats. The lack of effect of CORT in some cases could be explained by the fact that we received our rats from different vendors. We did see our expected effect of CORT, but only in male rats from Charles River and female rats. In the rats that we did see an increase in sign-tracking following CORT administration, we speculate that CORT exerts this effect by influencing dopamine activity in the nucleus accumbens (NAc), which, in turn, influences incentive learning. In support, systemic CORT administration has been shown to increase dopamine levels within the NAc (Piazza et al., 1996), and sign-tracking behavior is known to be dependent on NAc dopamine activity (Flagel et al., 2011; Saunders and Robinson, 2012; Iglesias et al., 2023).

The vendor effects we observed may arise from differences in how the rats from each vendor were bred and raised. Indeed, it is possible that genetic and/or environmental differences between vendors contributed to the effects we observed. We have previously reported effects of vendor on the propensity to sign-track and identified marked population structure among colonies from a given vendor, suggesting that different colonies are genetically distinct populations (Fitzpatrick et al., 2013). Thus, it is possible that random genetic drift or population bottlenecks reduce phenotypic heterogeneity and/or skew inherent behavioral tendencies. With respect to possible environmental effects, exposure to stress early in life has been shown to increase sign-tracking behavior in adulthood (Lomanowska et al., 2011). Furthermore, it is known that altered brain development resulting from early life stress affects an individual’s response to drugs of abuse and to stress in adulthood (Sinha, 2001; Ladd et al., 2004; Chen and Baram, 2016). In particular, exposure to stress during early life causes altered hippocampal glucocorticoid receptor (Type I and II) expression in adulthood, which affects the physiological response of corticosterone to stress (Ladd et al., 2004). It has also been shown that early life stress results in an increase in stress-induced dopamine transmission later in life (Meaney et al., 2002). Taken together, these findings suggest that early life stress may alter glucocorticoid activity, dopamine activity, and the way in which glucocorticoids and dopamine interact to modulate behavior (Lopez and Flagel, 2021). Thus, it is possible that rats from different vendors have experienced different levels of early life stress, which could in turn affect their tendency to sign-track and the impact of CORT in adulthood. Similarly, given different means of transportation to the lab, it is possible that the rats from different vendors experienced varying degrees of stress in adulthood, which is also known to affect sign-tracking behavior (Fitzpatrick et al., 2019). Future studies should assess hippocampal glucocorticoid receptor expression and baseline and stress-induced dopamine transmission and CORT levels in male and female rats from different vendors.

In addition to the vendor effects, interesting sex differences were revealed in this study. Although males and females were run separately in Experiments 1 and 2, our findings suggest that the influence of vendor on the effects of CORT during acquisition are different in males and females. CORT enhanced vendor differences in the acquisition of sign-tracking in males but enhanced the acquisition of sign-tracking in females, regardless of vendor. This difference may be explained by an interaction between CORT administration during acquisition and the inherent tendencies towards sign-tracking in these groups. In support, in Experiment 3, where male and female rats were run concurrently, female rats displayed significantly greater sign-tracking behaviors compared to males during the acquisition phase. In accordance with this finding, previous literature suggests that the female population tends to be skewed more towards sign-tracking relative to males (Hughson et al., 2019). Moreover, rats from Charles River displayed a greater tendency towards sign-tracking than those from Taconic with this vendor difference present in both sexes. Importantly, the groups that were shown to exhibit inherent tendencies towards sign-tracking in the acquisition phase of Experiment 3—females and rats from Charles River—are the same groups where CORT administration enhanced the acquisition of sign-tracking in Experiments 1 and 2. As discussed above, potential differences in glucocorticoid receptors, CORT levels, and dopamine transmission may contribute to vendor and sex differences in inherent tendencies towards sign-tracking and the ability of CORT to enhance these tendencies. In support, previous literature shows that female rats have higher basal and stress-induced plasma CORT levels (Critchlow et al., 1963; Kitay, 1961), and a more recent report suggests that sex differences in dopamine release in the NAc depend on strain (Rivera-Garcia et al., 2020). More work is needed to examine the role of potential sex and vendor differences in the underlying biological mechanisms that contribute to inherent tendencies to sign-track.

The current study revealed notable differences in CORT’s influence during acquisition vs. expression of sign-tracking. CORT treatment enhanced acquisition of sign-tracking in females and rats from Charles River in Experiments 1 and 2; however, once the conditioned response was acquired in Experiment 3, CORT administration failed to significantly enhance the expression of sign-tracking in these groups. One potential explanation is a ceiling effect; perhaps administering CORT to these rats would not make a measurable difference because they are hitting a ceiling in the expression of sign-tracking behavior. Another potential explanation is that CORT may influence underlying mechanisms important for the acquisition, but not expression, of sign-tracking behavior. As mentioned earlier, we speculate that CORT enhances the acquisition of sign-tracking through increasing dopamine transmission in the nucleus accumbens. Perhaps CORT administration during the expression phase did not significantly influence sign-tracking because the influence of dopamine signaling, particularly through D1 receptors, in the nucleus accumbens on sign-tracking behavior diminishes with more training (Clark et al., 2013). Further investigations are needed to directly test this hypothesis and to examine the neural mechanisms underlying the effects of CORT on sign-tracking behavior.

Conditioned reinforcement testing was used in our studies to gauge the incentive motivational value of the lever-cue as a function of prior treatment. Thus, regardless of treatment group, rats did not receive CORT treatment immediately prior to this test. In the groups with a heightened propensity to sign-track during PavCA (i.e., females and rats from Charles River), the lever-cue served as a more effective conditioned reinforcer. This was especially evident in Experiment 3, likely due to the rats experiencing more PavCA sessions (i.e., nine) than the other experiments (i.e., five) prior to the test. While systemic CORT treatment enhanced the acquisition (but not expression) of approach behavior toward the lever-cue, it did so without carry-over effects influencing the ability of the lever-cue to act as a conditioned reinforcer. Having CORT onboard during the conditioned reinforcement test may enhance the conditioned reinforcing properties of the lever-cue as it enhanced the inherent tendencies to attribute incentive value to the lever-cue during PavCA, although this would need to be tested directly.

These findings demonstrate that systemic administration of CORT can influence the propensity to attribute incentive value to reward cues by interacting with inherent tendencies. The effects of systemic CORT administration on the acquisition of PavCA behavior differs between male and female rats and those from different vendors. These findings underscore the importance of reporting vendor in scientific research and including vendor as a critical variable of analysis. The sex-dependent and vendor-dependent effects of CORT found in these studies capture the complexity of glucocorticoids, which are known to play a role in psychiatric disorders (McEwen & Akil, 2020; Piazza & Le Moal, 1996). We speculate that these differences are dependent on inherent tendencies towards sign- or goal-tracking behaviors based on sex and vendor, and perhaps the associated genetic and environmental conditions. Future studies should further examine these sex and vendor differences by assessing differences in baseline CORT levels and glucocorticoid receptor expression between sexes and vendors. In addition, we speculate that in rats where CORT promotes the acquisition of sign-tracking, it does so by influencing dopamine transmission in the NAc to enhance incentive learning. Future investigations are necessary to directly test this proposed mechanism.

## Acknowledgements

We would like to thank Charlotte Yang, Victor Sanchez Franco, and Jasmine Bhatti for technical assistance with portions of these experiments.

## Notes

### Competing Interest Statement

The authors have declared no competing interest.

